# Single Cell RNA-sequencing of BCG naïve and recurrent non-muscle invasive bladder cancer reveals a CD6/ALCAM-mediated immune-suppressive pathway

**DOI:** 10.1101/2025.02.13.638074

**Authors:** Ivan Juric, Emily E. Fink, Hong Qiu, Pierre-Emmanuel Desprez, Arvind Ravi, Mark Holton, Vladimir Makarov, Nima Almassi, Booki Min, Gad Getz, Timothy A. Chan, Tyler Alban, Angela H. Ting, Byron H. Lee

## Abstract

Non-muscle invasive bladder cancer (NMIBC) represents 70–80% patients with newly diagnosed bladder cancer, and Bacillus Calmette-Guérin (BCG) remains a cornerstone treatment for intermediate-and high-risk NMIBC to prevent disease recurrence and progression. However, many patients experience recurrence after induction BCG, posing significant challenges in the management of the disease. We conducted single cell RNA sequencing on freshly collected NMIBC samples, distinguishing between those naïve to BCG treatment and those that recurred post-BCG treatment. We observed a clear activation of inflammatory pathways across cell types during recurrence, but these were not associated with canonical immune checkpoint or T cell exhaustion phenotypes. Analysis of cell-to-cell communication revealed enhanced interactions between T cells and urothelial cells in BCG-recurrence, predominantly modulated by CD6 and ALCAM. Furthermore, we found CD6^hi^ T cells to be immunosuppressed and enriched in recurrent samples, suggesting a potential role for CD6 as an immune evasion signal in NMIBC.

**SIGNIFICANCE:** These findings uncover a novel mechanism responsible for bladder cancer recurrence after BCG treatment. Enhanced T cell-urothelial cell communication in recurrent tumors mediated by CD6 and ALCAM leads to an immunosuppressed state. Thus, CD6 may have potential as a therapeutic target to augment BCG response in non-muscle invasive bladder cancer.

## INTRODUCTION

Bladder cancer remains a significant global health burden, with non-muscle invasive bladder cancer (NMIBC) comprising the majority of cases at initial diagnosis. Intravesical Bacillus Calmette-Guérin (BCG) immunotherapy represents the gold standard for preventing recurrence and progression in intermediate- to high-risk NMIBC patients. However, up to 50% of patients will recur after their first induction BCG course (1–3). Moreover, approximately 30% of NMIBC patients will progress to muscle invasive bladder cancer, which is associated with high rates of occult metastatic disease.

The mechanisms underlying BCG resistance are complex and multifactorial, involving interactions between the tumor microenvironment, host immune response, and tumor intrinsic factors. Understanding the molecular basis of BCG resistance is crucial for the development of more effective therapeutic strategies and personalized treatment approaches in NMIBC, especially in a time of widespread BCG shortage (2,4). Multiple studies have sought to understand genomic and transcriptomic drivers of recurrence, with some success in identifying subtypes by mutation (5), copy number (6), and transcriptional signatures (7). However, these bulk genomic and transcriptomic analyses cannot fully uncover mechanisms of BCG resistance related to alterations in tumor-immune cell communication. To identify molecular factors associated with recurrence after BCG therapy, we employed single cell RNA sequencing (scRNA-seq) to analyze the heterogeneity of tumor cell populations and their interactions with the immune microenvironment. This study compared NMIBC samples obtained prior to BCG treatment with those that recurred after BCG therapy.

## RESULTS

### Cell type composition of BCG naïve and recurrent tumors

To understand the genomic and molecular drivers of BCG resistance, we implemented a protocol for rapid dissociation and scRNA-seq from cystoscopically resected NMIBC specimens along with parallel whole exome sequencing. Our final cohort consisted of 27 samples from 23 patients, with 14 samples collected prior to BCG treatment (“naïve” samples; n=13 patients) and 13 samples collected following BCG treatment (“recurrent” samples; n=12 patients; Table 1, Supplementary Fig. S1). The median follow-up for the naïve group without recurrence was 37.7 months (range 27.6 – 43.3 months). For the recurrent group, the median time to event was 5.8 months (range 1.8 – 13.2 months) from last BCG instillation. Within our BCG-naïve group, 3 of 13 patients eventually recurred. One patient was found to have muscle invasive bladder cancer on restaging transurethral resection and was not treated with BCG. In the 2 other cases, we were able to obtain matched naïve and recurrent specimens, enabling limited longitudinal analysis. Initial somatic profiling of our study cohort (n=23) revealed strong similarities between BCG naïve and recurrent tumors, including known mutations in *FGFR3*, *PIK3CA*, *KMT2D*, *KDM6A*, *KMT2C*, and *ARID1A* (Fig. 1A) (8). *CDKN2A* loss was frequently identified, affecting 70% of all tumors, and mutational signature analysis showed known signatures of NMIBC including APOBEC (SBS13 and SBS2) (Fig. 1A). Looking further into the genomic alterations, we searched for copy number changes that might be driving recurrence and noted no significant changes between naïve and recurrent samples (all q-values > 0.05, GISTIC); however, commonly reported NMIBC alterations like gain of chromosome 1q and loss of chromosome 9p and 9q were evident (Fig. 1B).

**Figure 1.**
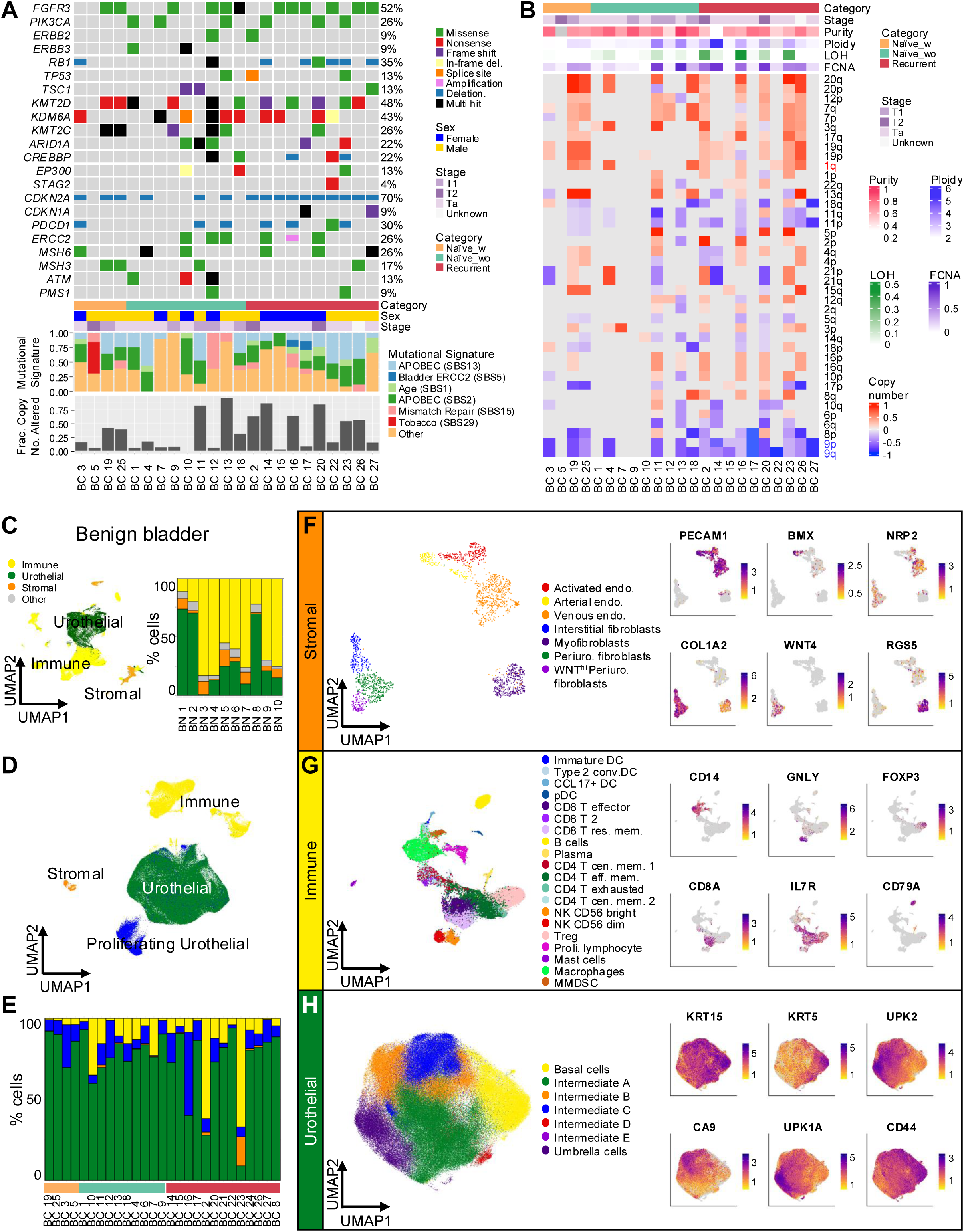
Genomic and single cell transcriptional profiling of NMIBC. A) Oncoplot summary of top NMIBC related genomic alterations separated by two primary groups of interest, BCG naïve (orange and green) and BCG-treated samples acquired after recurrence (recurrent, red). Clinical follow-up further identified two groups within the BCG naïve cohort, those patients who were later treated with BCG but did not recur at the time of follow up (naïve_wo, green), and those who recured post-BCG (naïve_w, orange). B) Summary of copy number alterations across chromosome arms per sample. Common gain (1q) and loss (9p and 9q) regions for bladder cancer are labeled with Red/Blue row names. Fraction copy number alterations, loss of heterozygosity, tumor purity, and tumor ploidy are also presented above copy number alteration heatmap with legend to the right for each unit. C) UMAP containing 229,558 cells from 10 benign bladder samples partitioned into three major compartments for reference mapping of NMIBC samples (stromal, immune, and urothelial). Suspected doublets were classified as other. The proportions of each cell class in an individual sample are summarized in a histogram (right). D) UMAP containing 250,229 cells from 27 bladder cancer samples partitioned into stromal, immune, urothelial, and proliferating urothelial compartments using the benign bladder as a reference. E) Fraction of major cell compartments across NMIBC samples annotated with their BCG treatment status. F-H) Re-clustering and detailed naming of the stromal (F), immune (G), and urothelial (H) compartments within NMIBC samples along with representative feature plots (legend colors = average expression).

**Table 1.**
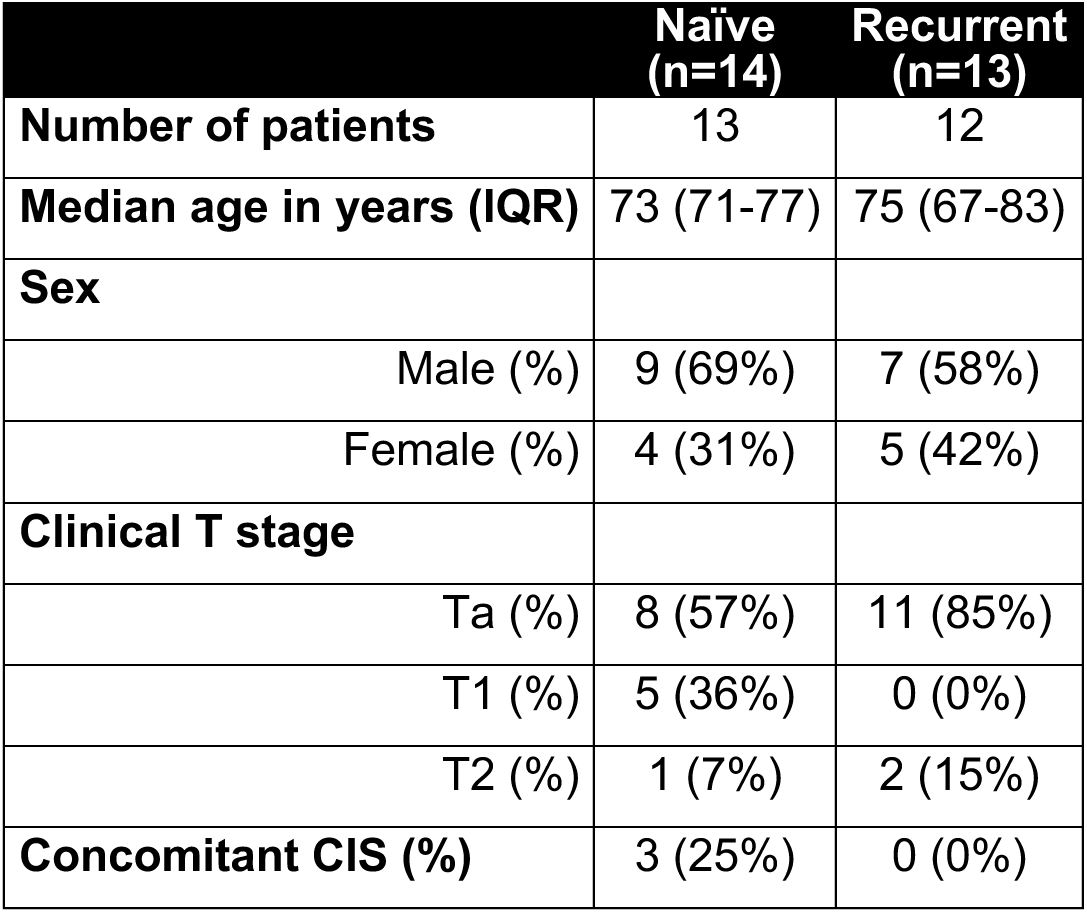
Clinical cohort characteristics.

Given the absence of dramatic differences in somatic events between BCG naïve and recurrent samples – our sample size notwithstanding – we hypothesized that disease recurrence may be transcriptionally regulated, and that single cell RNA-sequencing (scRNA-seq) may help elucidate the cellular state changes underlying BCG resistance. As NMIBC cells may express aberrant transcriptional profiles depending on their degree of differentiation, we performed scRNA-seq on n=10 freshly isolated bladder biopsies away from tumor with normal appearing urothelium and used these samples as a reference atlas to assist with cell type annotation (Fig. 1C, Supplementary Table S1). Within these benign bladder samples, most had predominantly immune or urothelial compartments, with smaller contributions from stromal cell types (Fig. 1C). Detailed naming of each compartment was performed (Supplementary Fig. S2), and then we mapped our cancer cell dataset to corresponding categories of Urothelial, Immune, and Stromal, along with a minor component of Proliferating Urothelial (Fig. 1D). Of these, the urothelial compartment was typically predominant, as expected by tumor sampling procedure which involves biopsy of their intraluminal aspect (Fig. 1E). Next, we isolated and re-clustered each compartment to better resolve individual cell types (Fig. 1F-H). Stromal cells showed expected fibroblast and endothelial cell subtypes, including myofibroblasts and WNT high periurothelial fibroblasts; however, given their limited representation in most samples, these cell populations were generally underpowered for more detailed analysis (Fig. 1E). For the immune compartment, a combination of reference mapping, AI-based naming, and expert curation were used to ensure rigorous classification of a total of 20 immune cell types across 27 samples (Supplementary Fig. S3). Lastly, naming of the urothelial compartment was performed by reference mapping to normal urothelial populations and assessment of basal cell and umbrella cell markers such as *KRT15*, *KRT5*, *UPK2*, and *UPK1A* (Fig. 1H). As expected with the variable differentiation state of NMIBC, we observed multiple clusters of intermediate cell types likely in transitory states between basal and umbrella cell identities (Fig.1H).

### Interferon signaling is broadly activated across cell types after BCG treatment

After harmonization of single cell populations across patients, we sought to identify changes in cell-type abundance that might exist between naïve (n=15) and recurrent samples (n=13). When represented as a percentage of their respective compartments, we saw no statistically significant differences in cell type abundance between the naïve and recurrent specimens, with or without FDR correction (Fig. 2A). Without strong evidence of compositional changes after BCG treatment, we reasoned that BCG resistance may be driven by more nuanced changes in the cellular state of individual sub-populations. To resolve these changes in cell state, we performed cell-type specific differential expression to identify genes enriched in recurrent samples (n=15) relative to naïve samples (n=15) (Supplementary Table S2). Gene set enrichment analysis of these genes using the Hallmark pathways showed surprisingly broad activation of inflammation-related pathways across compartments, such as tumor necrosis factor alpha (TNFA) signaling, interferon gamma (IFNG) signaling, and interferon alpha (IFNA) signaling (Fig. 2B, Supplementary Table S3). These results suggest that the increased inflammation described in bulk RNA-seq studies of BCG resistance may reflect contributions from a relatively broad collection of distinct cell types.

**Figure 2.**
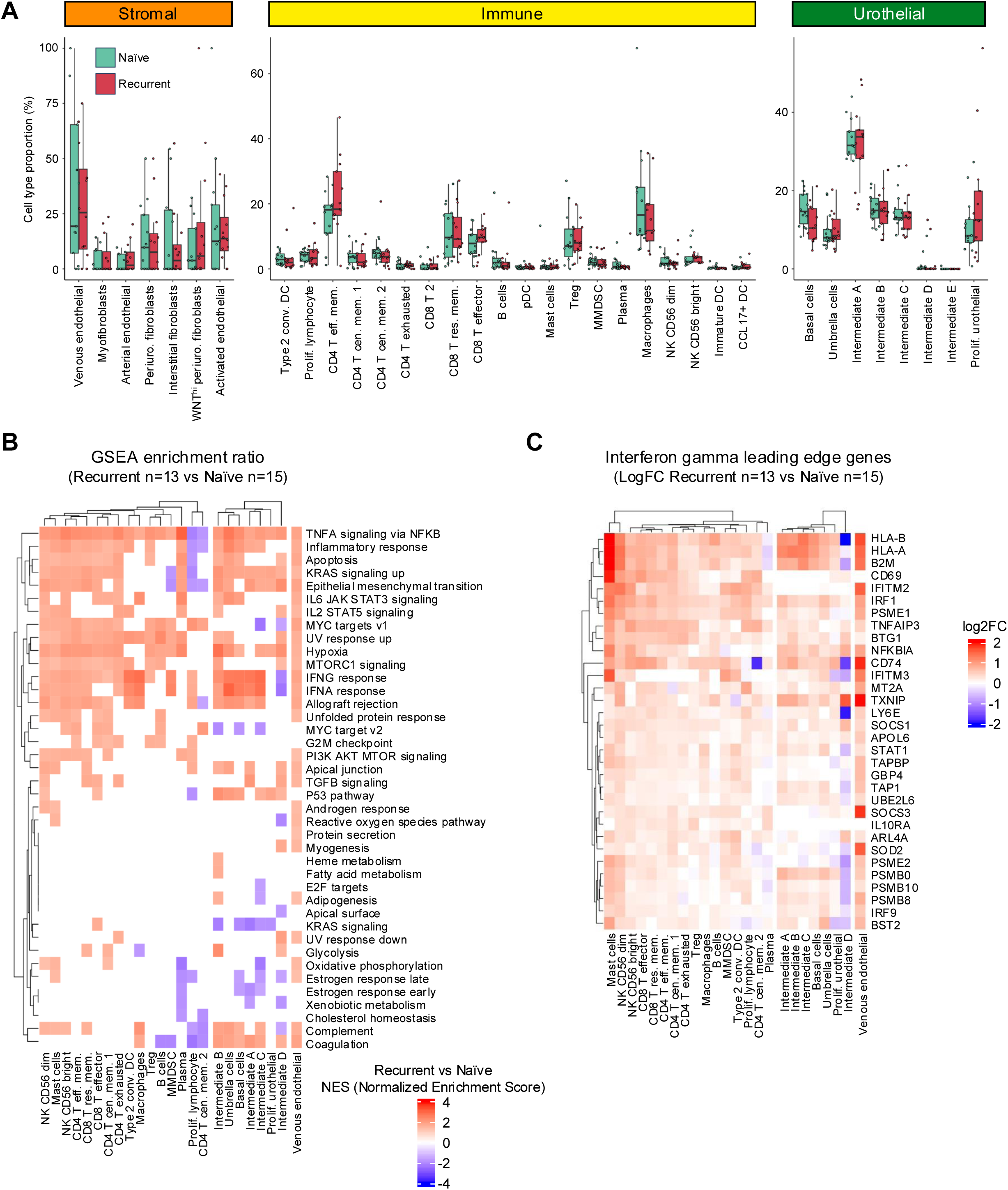
Differential cell abundance and differential gene expression between BCG-naïve and BCG-recurrent NMIBC. A) Differential abundance analysis was performed using the proportion of cells per sample comparing naïve and recurrent samples (Wilcoxon test, no p-values met significance threshold of p<0.05). B) Heatmap of enriched hallmark pathways from GSEA analysis of differentially expressed genes per cell type between naïve and recurrent samples. The color scale represents the normalized enrichment score (NES) which met p-value threshold of p<0.05. Non-significant NES scores were not plotted (white). Positive NES values (red) indicate pathway enrichment in recurrent samples. C) Log2 fold change of leading-edge genes for the interferon gamma (IFNG) pathway (red= enriched in recurrent, blue= enriched in naïve

To further understand the genes driving each immune activation signature, we isolated the leading-edge genes from the IFNG pathway and saw a common activation of antigen presentation genes such as *HLA-A*, *HLA-B*, and *B2M* (Fig. 2C, LogFC comparing recurrent n=13 and naïve n=15). Many of these genes overlapped with leading-edge genes of the IFNA pathway (Supplementary Fig. S4A, LogFC comparing recurrent n=13 and naïve n=15), while those in the TNFA pathway appeared to be more closely related to cytokine response and proliferation such as *FOS*, *JUNB*, *JUN*, and *CD69* (Supplementary Fig. S4B). To further resolve how specific the IFNG activation pathway is to recurrence, we utilized our 2 patients which had matched samples from the BCG-naïve and recurrent settings. For patient 1, who had one naïve sample and two recurrent samples, we observed a parallel increase in interferon pathway gene expression – based on the log2 ratio of IFNG signature genes relative to the naïve baseline – observed at the cohort-wide level (Supplementary Fig. S4C). Similarly, in patient 2, who had two naïve samples and one recurrent sample, we saw relatively lower IFNG pathway expression prior to treatment (Supplementary Fig. S4D). Taken together, observations from both unmatched and matched samples show that BCG-recurrence is enriched for immune activation signatures of the IFNG pathway across all cellular compartments.

### CD6/ALCAM interactions are increased in BCG recurrent samples and associated with T cell inhibition

While pathway enrichment of BCG recurrent samples vs Naive identified increased inflammatory signatures during recurrence, it was unclear how NMIBC evades this enhanced immune activation. To surface more nuanced mechanisms of immune resistance related to specific cellular interactions in the microenvironment, we performed cell-cell communication analyses using CellChat (9). For an initial overview of interaction changes, we grouped cell types into major categories, Urothelial, Stromal, T cells, B cells, NK cells, Myeloid cells, and Dendritic cells. Within these major groups, we summarized the overall cell-cell interactions by comparing communication patterns in naïve and recurrent samples. We found that recurrent samples (n=15) had increased Urothelial-to-Urothelial (n=451), Urothelial-to-T cells (n=175), and Stromal-to-Urothelial (n=177) interactions (Fig. 3A). We next focused specifically on outgoing and incoming interactions between T cells and urothelial cells to precisely understand potential alterations in immune recognition and clearance. Across all comparisons in the naïve samples (n=13), we saw enrichment for typical immune trafficking and surveillance interactions such as those between MHC class II genes and CD4 as well as a variety of CD44 ligands and CD44, a marker of basal bladder cancer cells that correlates with recurrence, invasion, and poor prognosis (10) (Supplementary Fig. S5A). There was also evidence of some immune suppressive interactions between T cells and urothelial cells like TIGIT and NECTIN2. Within the recurrent tumors (n=15), we saw high enrichment of ADGRE5 on T cells interacting with CD55 on urothelial cells (11), a pathway which has been shown to be associated with tumor progression and stemness across various cancer types. Increased communication between urothelial MDK and T-cell NCL was also identified in the recurrent tumors, and this intercellular signaling pathway has been associated with immunosuppressed populations that are PD1 high in other cancers such as triple negative breast cancer (12). Despite seeing some mild enrichment of immune exhaustion related pathways, we saw no major expression changes in well described immune checkpoints (e.g., PD1, PDL1, CTLA4, TIGIT) to explain recurrence after BCG treatment (Supplementary Fig. 5B).

**Figure 3.**
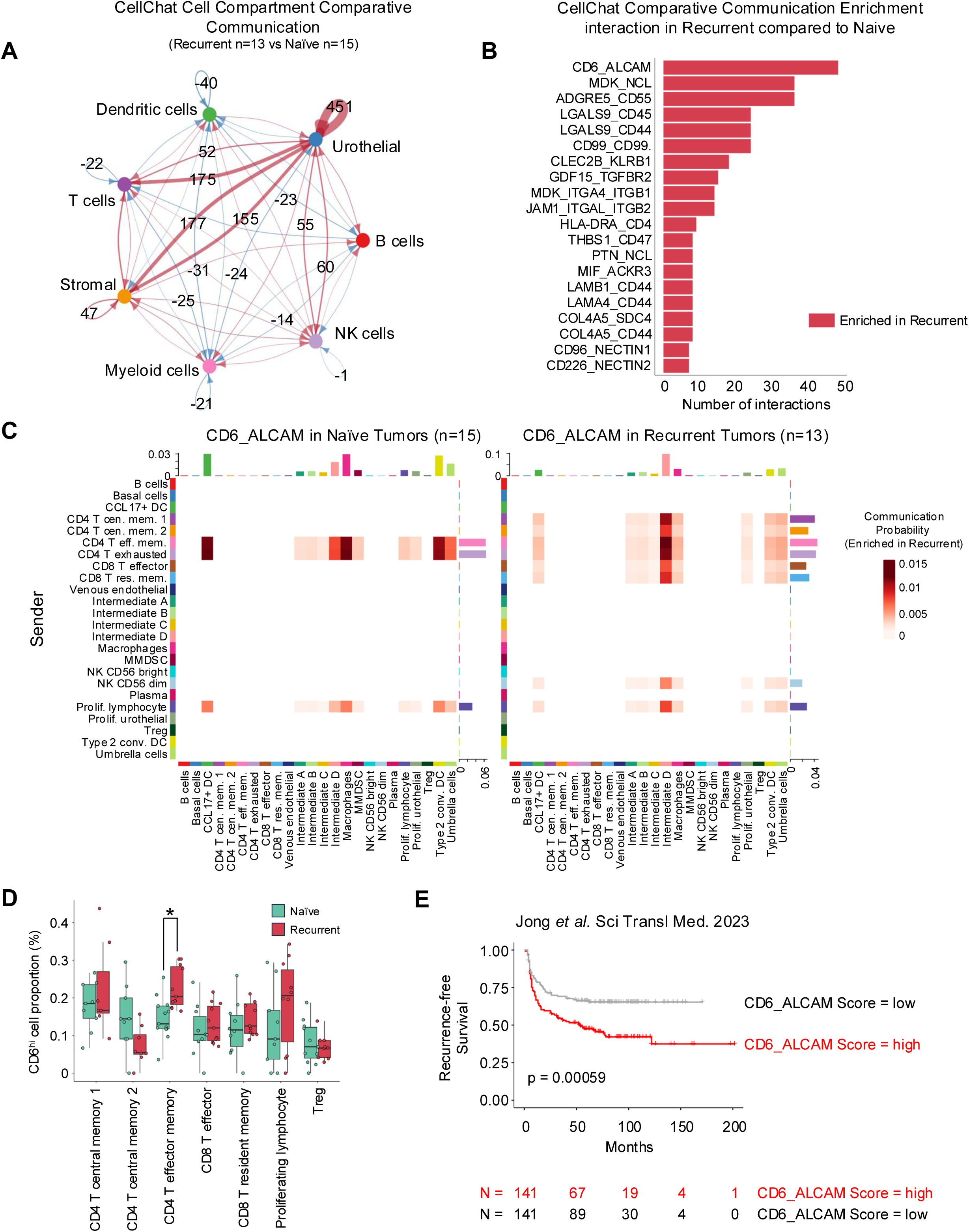
Comparative analysis of cell communication networks in BCG naïve and recurrent samples. A) The differences in the numbers of cell-to-cell interactions predicted by CellChat for naïve and recurrent samples are plotted. Red arrows indicate increased interactions in recurrent samples. Blue arrows indicate decreased interactions in recurrent samples. The thickness of the connecting lines relates to the number of differential interactions, with the largest changes in either direction labeled with exact numbers. B) Summary plot of enriched interactions between T cells and urothelial cells (aggregate of T cells-to-Urothelial and Urothelial-to-T cells interactions from Supplementary Fig. S4A) in recurrent samples compared to naïve. C) Heatmaps of CD6-ALCAM communication probability score between cell types in naïve (left) and recurrent (right) samples. D) Fraction of CD6^hi^ cells in each T cell population in naïve (green) and recurrence (red) samples. The asterisk indicates statistical significance (Mann Whitney U test, Benjamini-Hochberg-adjusted p=0.018). E) Time to disease progression post-BCG treatment for Jong *et al.* NMIBC cohort (n=282), stratified by median CD6/ALCAM dual gene signature score. P-value was derived from log-rank test.

To better gauge alterations in shared interactions between T cells and urothelial cells, we summed the total interactions that were enriched during recurrence across T cells-to-Urothelial and Urothelial-to-T cells (Fig. 3B). The most upregulated interaction was between CD6 on T cells and ALCAM (CD166) on urothelial cells. In the BCG-naïve samples, only a limited number of strong CD6 and ALCAM interactions were observed, linking CD4 T-cell populations to CCL17+ Dendritic cells and Type 2 conventional dendritic cells (left panel in Fig. 3C). In contrast, recurrent tumors exhibited a broader collection of T cells, including CD4 Central memory, CD8 effectors, and CD8 memory T cells, that interacted with urothelial cell types via CD6/ALCAM (right panel in Fig. 3C). NK cells (CD56 dim) were also identified as increased in CD6 interactions broadly across groups. Given the increase in CD6-mediated interactions in recurrence, we next sought to determine if the abundance of CD6 expressing T cells changed following recurrence. Classifying cells as CD6 high (CD6^hi^) vs. low (CD6^lo^) based on median expression, we compared naïve and recurrent samples based on the abundance of CD6^hi^ cells within each T cell subset (Fig. 3D). CD4 effector cells showed a significant increase in the proportion of CD6^hi^ cells following recurrence (Mann Whitney U test, Benjamini-Hochberg-adjusted p=0.018), with proliferating lymphocytes also showing a trend in this direction (Fig. 3D). In addition to changes in abundance, we assessed whether CD6^hi^ vs. CD6^lo^ cells may have differences in gene expression. Notably, CD6^hi^ cells demonstrated a broad loss of activation markers including *IL2RB*, *IRF1*, *IL2RG*, *TIGIT*, *TOX*, *JAK1*, *TOX4*, and *TNFAIP3* (Supplementary Fig. S5C).

To further understand how the identification of CD6/ALCAM from our single-cell interactions studies might help inform response to BCG, we identified a bulk RNA-seq study (7) with n=282 non-muscle invasive bladder tumors that included time to BCG recurrence. With the hypothesis that CD6/ALCAM high tumors are immunosuppressed via T cell inactivation and are more likely to recur after BCG treatment, we scored each sample for its expression of CD6, ALCAM, or CD6/ALCAM gene set for comparison (Supplementary Table S4). Using individual gene expression of CD6 (Supplementary Fig. S5D) or ALCAM (Supplementary Fig. S5E) with a median cutoff, we observed significantly shorter times to high grade recurrence post-BCG treatment (log rank test, p=0.014 and p=0.04 respectively). Moreover, when we stratify the cohort using the combination gene set of CD6/ALCAM, an even stronger effect emerged showing CD6/ALCAM high samples with significantly shorter time to BCG recurrence than CD6/ALCAM low samples (Fig. 3E, log rank test, *p*=0.00059).

## DISCUSSION

In this study, we aimed to elucidate the molecular drivers of recurrence after Bacillus Calmette-Guérin (BCG) treatment in non-muscle invasive bladder cancer (NMIBC) through a comprehensive scRNA-seq analysis of freshly isolated tumor specimens collected before and after BCG treatment. In a time of widespread BCG shortage, identifying mechanisms responsible for disease recurrence has become even more critical, so that we can devise therapeutic strategies to augment BCG efficacy or judiciously offer clinical trial enrollment. While there have been multiple attempts to identify mechanisms of recurrence using bulk RNA-seq (7,13) and liquid biopsies (14,15), we used single cell analysis of BCG-naïve and recurrent NMIBC samples to elucidate potential differences in cell proportion, cell state, and intercellular communication. Upon harmonization of single-cell populations across patients, we identified broad activation of inflammation-related pathways in BCG recurrent samples, particularly interferon gamma (IFNG) signaling, across multiple cellular compartments. The interferon signaling pathway and antigen presentation machinery were both upregulated, suggesting a heightened immune response associated with exposure to BCG. Longitudinal analysis of matched samples further supported the consistent immune activation and interferon type II response in BCG-treated cancers. While this seems counter intuitive to see increased inflammation in recurrent tumors, a prior bulk RNA-seq study has also identified increased inflammatory signatures, including IFNG, IL-2, and MHC class I genes, to correlate with BCG resistant samples (7). The authors further defined this subset as BRS3 subtype and hypothesized that immune exhaustion may explain their lack of response; but they did not see striking signals of standard immune checkpoints like PD-1 (7). Due to the limitations of bulk RNA-seq, the authors could not further resolve populations to understand how these inflamed tumors remained resistant to BCG (7), however here we expand on the mechanisms of immune evasion during this broad inflammatory state.

While gene expression and pathway analysis all indicated increased immune activation, we performed in-depth cell-to-cell communication analysis, which revealed increased interactions between T cells and urothelial cells in recurrent samples, particularly through the CD6/ALCAM axis. We further identified that broadly, T cells with high CD6 expression had a reduced activation capacity. CD6 is a transmembrane glycoprotein primarily expressed on the surface of T cells, where it plays a crucial role in modulating T cell activation and function, particularly in the context of immune responses and tolerance (16–18). CD6 interacts with multiple ligands including ALCAM (CD166), CD44, and CDCP1 (CD318), primarily through its extracellular domain and signals through its cytoplasmic tail via modulation of Ras (19). Importantly, studies have shown that mutation of the intracellular signaling domain of CD6 results in enhanced activation capacity i.e. CD6 WT signaling reduces immune activation (19). The CD6/ALCAM interaction is thought to be involved in signaling as well as T cell adhesion to antigen-presenting cells (APCs) and the formation of immunological synapses, thereby regulating T cell activation and immune responses. The dysregulation of CD6-mediated signaling has been associated with autoimmune diseases, including rheumatoid arthritis, multiple sclerosis, and systemic lupus erythematosus (20–23). CD6 has been proposed as a potential immunotherapeutic target for breast cancer, lung cancer, and prostate cancer (24), and CD6^+^/CD8^+^/PD-1^+^ T cells have been shown to be deficient in Granzyme, perforin, and interferon gamma production compared to CD6^-^/CD8^+^/PD-1^+^ T-cells (25).

Validation of the CD6/ALCAM signature was performed using bulk RNA-seq, where we found that samples expressing high levels of both CD6 and ALCAM were present prior to BCG treatment and strongly correlated with overall response to BCG. This indicates there is likely a pre-programing of T cells that has already occurred prior to BCG treatment, similar to the findings of others that an inflamed state prior to BCG treatment may indicate a poor outcome (7). While this finding may seem counterintuitive compared to other cancer types, where an immune “hot” tumor microenvironment indicates favorable outcomes, it is important to note that BCG treatment is rather unique in the use of a live bacteria as a therapeutic agent that activates an innate response followed by induction of adaptive immunity (26). Based on our scRNA-seq results, validation in bulk RNA-seq data, and previous studies showing CD6 as an immune response regulator, we believe that targeting CD6 can potentially reduce recurrence after BCG treatment (27).

Overall, our findings shed light on the immune cell interactions potentially driving BCG resistance and highlight a need to further investigate the role of CD6/ALCAM interaction within NMIBC. While we are *in silico* estimating cellular interactions via CellChat (9), the bulk RNA-seq analysis strongly indicates a role for this signaling cascade’s involvement in mediating recurrence after BCG treatment. Furthermore, other interactions that were identified have been shown to be present in NMIBC and/or have been sought after targets such as MDK/NCL (28,29) and TIGIT/NECTIN2 (30,31). Future studies should seek to understand how the interaction drives immune suppression in NMIBC, and exploration of clinical interventions targeting CD6/ALCAM may be warranted to improve treatment outcomes for patients prior to or in combination with BCG.

## METHODS

### Patients, Sample Collection, Treatment, and Follow-up

Patients with a new bladder tumor diagnosed by axial imaging and/or cystoscopy were consented for specimen collection and molecular analysis under and Institutional Review Board-approved protocol adhering to U.S. Common Rule guidelines at Cleveland Clinic. Tumors were identified with a cystoscope, and a cold cup forceps was used to biopsy the intra-luminal portion. Control specimens were biopsied in a similar fashion from areas with normal appearing urothelium at least 5 cm away from any tumor site or suspicious lesion. Tissue pieces were either snap frozen immediately in liquid nitrogen or dissociated for 10x Genomics 3’ single cell RNA-seq library preparation as previously described (32). All patients with T1 HG or multifocal Ta HG disease underwent repeat transurethral resection prior to BCG treatment. All patients received six instillations of BCG (TICE), and surveillance was performed every 3 months with white light cystoscopy and urine cytology.

### Whole Exome Sequencing

Genomic DNA was extracted from frozen tumor tissues and quantified using nanodrop. The sequencing library was made using SureSelectV5 library kit following manufacturer’s protocol and sequenced using paired end 150 bp reads to an average coverage of 187x (range 151x-263x). The sequencing reads were aligned to GRCh38 using STAR. A combination of five different mutation callers (Mutect2, SomaticSniper 1.0.4, VarScan V2.4.3, Strelka v2.9.10 and Platypus 0.8.1) were used to identify SNVs. Small insertions and deletions (indels) were determined using Mutect2, Varscan v2.4.3, Strelka v2.9.10, and Platypus 0.8.1. We used the SnpEff variant annotation and effect prediction tool to determine effect of called variants. Our variant reporting criteria are as follows:

- Tcov > 10 & Taf ≥ 0.04 & Ncov > 7 & Naf ≤ 0.01 & Tac > 4 were set to pass.
- Common SNPs were eliminated by comparison to snp142.vcf.
- Rare variants found in dbSNP were kept if Naf = 0.
- Variants with Tcov < 20 or Tac < 4 were marked as low confidence.
- Only Variants called by more than 1 caller were reported.

Common variants in gnomAD v 2.1.1 (March 2019 release; gnomAD also incorporates Exome Aggregation Consortium (ExAC)) were excluded. Additional optimization and filtering were applied for insertions/deletions (INDELS). INDELS in blacklisted regions (https://www.encodeproject.org/annotations/ENCSR636HFF/) and low mappability regions were excluded.

Mutation load was determined as the total number of non-synonymous mutations passing filters. Previously described signatures of mutational processes were determined in each sample using non-negative least-squares regression as provided by the R package deconstructSigs v1.8.0 using the COSMIC signatures as the mutational signature matrix. Mutational profiles were analyzed from MAF files using R package maftools version 2.12.0. Gistic2.0 was used for analyzing copy number change from facets segment values.

### Single Cell RNA-seq mapping and cell naming

Fastq files were mapped to the GRCh38 reference human genome using Cellranger (v5.0.0). Cells containing less than 600 genes and/or more than 30% mitochondrial and ribosomal genes were removed. Sample-specific Seurat objects were created using Seurat (v4.3.0), then normalized using Seurat’s SCTransform. Samples were then split into benign bladder (10 samples) and cancer (27 samples) sets. Each set was integrated independently, based on variable features using Seurat’s IntegrateData function. Cells from the benign bladder set were then divided into Immune, Stromal, and Urothelial compartments, re-clustered, and named based on marker genes. We also identified and named clusters containing doublets (cell clusters that contain cells with abnormal number of genes or markers characteristic for multiple different cell types).

To help name cancer cells, we used Seurat’s label transfer method (33) to map cancer set to benign bladder set. Provisional names were then used to identify Immune, Stromal, and Urothelial compartments. Proliferating Urothelial cells were assigned to separate compartment based on the presence of cell proliferation markers. Low quality cells in the Urothelial compartment were further filtered out by removing all cells that contained less than 2000 nFeatures per cell. Cells in each compartment were then named in an iterative process that involved manually examining cell markers for each cluster, naming clusters that consisted of a single cell type, and re-clustering the remaining cells using Harmony (v0.1.0).

### Differential gene expression, differential abundance, GSEA, and CellChat analyses

Differential gene expression was assessed with Seurat’s FindMarkers function using a negative binomial generalized linear model (test_use = ‘negbinom’) with patient as a covariate to account for several samples that came from the same individual. Parameter logfc_threshold was set to 0.05. Other parameters were set to default. Differential expression (DE) analysis was performed only for comparisons in which there were more than 100 cells present for analysis in recurrence and naïve groups.

The output of DE analysis was the input for gene set enrichment analysis, performed using the GSEA function from clusterProfiler (v4.4.4) and HALLMARK gene sets. All genes, regardless of p-value, were sorted from highest to lowest log2 fold change and used for the analysis.

Longitudinal analysis of two individuals with tumors taken at multiple time points were used to investigate the change in IFNG related gene sets using ssGSEA scores. Differential abundance analysis was performed by comparing fractions of different cells between different groups using Wilcoxon rank sum test as implemented in R. Cell types with more than 100 cells per group were considered for this analysis. Heat maps were generated using ComplexHeatmap and circlize packages. To group cells into CD6 (ALCAM) high and CD6 (ALCAM) low groups, we first split cells into those that express the gene (count > 0) and those that do not and filter out all cell types for which less than 100 cells expressed the gene. Next, we found the mean log-normalized gene expression of gene-expressing cells and used that value to separate low from high groups. Cell-to-cell interaction was performed using CellChat 1.6.1 (9). Only cell types present in at least 5 samples and with >100 total cells were included in the comparisons. We summed the significant ligand-receptor pairs for naïve and recurrent samples between Urothelial cells, Urothelial cells and T cells, and Stromal cells and Urothelial cells. Then, we subtracted the naïve sum from the recurrent sum to identify alterations in cellular communications between groups.

### Bulk RNA-seq analysis

Validation of CD6/ALCAM signature association with BCG response was performed using the bulk RNA-seq data from Jong *et al.* (7), where gene expression data for primary samples (n=282) was extracted from Supplemental Data Files S1 and combined with metadata from Supplemental Table S2. For single gene analysis of CD6 and ALCAM, the median expression value was used to stratify high/low groups. Recurrence free survival analysis was performed using time to progression (“Time_to_HG_recur_or_FUend”) and the censor ("HG_recur_BCG_failure”) with log-rank method in R package survminer. Dual signature of CD6/ALCAM was scored using the R package gsva with the ssGSEA method. The median ssGSEA score was then used to categorize samples into high/low groups for progression free survival analysis.

## Acknowledgements

We thank the patients treated at the Cleveland Clinic for donation of samples for this study and Dr. Colin Dinney for critically reading this manuscript.

## Data availability

Deposition of all raw whole exome sequencing data into dbGaP is in process. Single cell RNA-seq raw and processed data is available at GEO under accession number GSE269877.

---

**Supplementary Figure S1. Flow diagram of patients used for this study.**

**Supplementary Figure S2. Benign bladder scRNA-seq analysis.** A-C) Detailed cell typing in stromal (A), immune (B), and urothelial (C) compartments was performed and the fraction of cells within each compartment per sample are represented as cell abundance/sample composition bar graphs. D-F) Detailed cell naming was performed by using AI based prediction tools in BioTuring software along with manually curated markers as represented in dot plots of key markers.

**Supplementary Figure S3. NMIBC sample scRNA-seq analysis.** A-C) Detailed cell typing in stromal (A), immune (B), and urothelial (C) compartments was performed and the fraction of cells within each compartment per sample are represented as cell abundance/sample composition bar graphs. D-F) Detailed cell naming was performed by using AI based prediction tools, mapping to benign bladder reference UMAP, and validated using manually curated markers as represented in dot plots of key markers.

**Supplementary Figure S4. Inflammatory pathway enrichment analysis.** A) Log2 fold change of leading-edge genes for the interferon alpha (IFNA) pathway (red= enriched in recurrent, blue= enriched in naïve). B) Log2 fold change of leading-edge genes for the TNF alpha pathway (red= enriched in recurrent, blue= enriched in naïve). C) Patient 1 had a single sample prior to BCG treatment (naïve, orange) and two samples after BCG recurrence (red). Expression of IFNG pathway genes were summarized as ssGSEA scores for each cell type with >100 cells. Using the BCG naïve sample (BC_3) as a reference, the log2 ratio change in IFNG ssGSEA score was plotted for each cell type for the BCG recurrence samples (BC_20 and BC_14). Positive log2 ratio (red) indicates increases in recurrence, and negative ratio (blue) indicates decrease in recurrence. D) Patient 2 had two samples prior to BCG treatment (naïve, orange) and a single sample after BCG recurrence (red). Expression of IFNG pathway genes were summarized as ssGSEA scores for each cell type with >100 cells. Using the BCG recurrence sample (BC_26) as a reference, the log2 ratio change in IFNG ssGSEA score was plotted for each cell type for the BCG naïve samples (BC_19 and BC_25). Positive log2 ratio (red) indicates decrease in recurrence, and negative ratio (blue) indicates increase in recurrence.

**Supplementary Figure S5. T cell characterization in BCG recurrence.** A) Top enriched interactions between T cells-to-Urothelial, T cells-to-T cells, Urothelial-to-T cells, and Urothelial-to-Urothelial in naïve (green) and recurrent (red) samples. B) Expression of immune checkpoint related gene in T cell populations with >100 cells per sample in naïve and recurrent specimens. C) Heatmap of differential gene expression in curated T cell activation genes (Nanostring T cell activation gene set) between CD6^hi^ and CD6^low^ (stratified by median CD6 expression) T cell populations. D-E) Time to disease progression post-BCG treatment for Jong *et al.* NMIBC cohort (n=282), stratified by median CD6 (D) and ALCAM (E) gene expression. P-values were derived from log-rank test.

**Supplementary Table S1.** Single cell RNA-seq quality control assessment using Cellranger metrics. Post alignment quality control metrics were collected using Cellranger 5.0.0 as described in the methods and recorded on a per sample basis.

**Supplementary Table S2.** Differential gene expression between naïve and recurrent samples for each cell type (each tab).

**Supplementary Table S3.** Hallmark pathway gene set enrichment analysis of differentially expressed genes between naïve and recurrent samples for each cell type (each tab).

**Supplementary Table S4.** Jong *et al.* bulk RNA-seq cohort data (from original publication supplementary materials) with ssGSEA scores for CD6, ALCAM, and CD6/ALCAM dual signatures.

## REFERENCES

1. Chamie K, Litwin MS, Bassett JC, Daskivich TJ, Lai J, Hanley JM, et al. Recurrence of high-risk bladder cancer: a population-based analysis. Cancer 2013;119(17):3219–27 doi 10.1002/cncr.28147.

2. Packiam VT, Werntz RP, Steinberg GD. Current Clinical Trials in Non-muscle-Invasive Bladder Cancer: Heightened Need in an Era of Chronic BCG Shortage. Curr Urol Rep 2019;20(12):84 doi 10.1007/s11934-019-0952-y.

3. van Rhijn BW, Burger M, Lotan Y, Solsona E, Stief CG, Sylvester RJ, et al. Recurrence and progression of disease in non-muscle-invasive bladder cancer: from epidemiology to treatment strategy. Eur Urol 2009;56(3):430–42 doi 10.1016/j.eururo.2009.06.028.

4. Bandari J, Maganty A, MacLeod LC, Davies BJ. Manufacturing and the Market: Rationalizing the Shortage of Bacillus Calmette-Guerin. Eur Urol Focus 2018;4(4):481–4 doi 10.1016/j.euf.2018.06.018.

5. Pietzak EJ, Bagrodia A, Cha EK, Drill EN, Iyer G, Isharwal S, et al. Next-generation Sequencing of Nonmuscle Invasive Bladder Cancer Reveals Potential Biomarkers and Rational Therapeutic Targets. Eur Urol 2017;72(6):952–9 doi 10.1016/j.eururo.2017.05.032.

6. Lindskrog SV, Prip F, Lamy P, Taber A, Groeneveld CS, Birkenkamp-Demtroder K, et al. An integrated multi-omics analysis identifies prognostic molecular subtypes of non-muscle-invasive bladder cancer. Nat Commun 2021;12(1):2301 doi 10.1038/s41467-021-22465-w.

7. de Jong FC, Laajala TD, Hoedemaeker RF, Jordan KR, van der Made ACJ, Boeve ER, et al. Non-muscle-invasive bladder cancer molecular subtypes predict differential response to intravesical Bacillus Calmette-Guerin. Sci Transl Med 2023;15(697):eabn4118 doi 10.1126/scitranslmed.abn4118.

8. Minoli M, Kiener M, Thalmann GN, Kruithof-de Julio M, Seiler R. Evolution of Urothelial Bladder Cancer in the Context of Molecular Classifications. Int J Mol Sci 2020;21(16) doi 10.3390/ijms21165670.

9. Jin S, Guerrero-Juarez CF, Zhang L, Chang I, Ramos R, Kuan CH, et al. Inference and analysis of cell-cell communication using CellChat. Nat Commun 2021;12(1):1088 doi 10.1038/s41467-021-21246-9.

10. Duex J, Theodorescu D. CD44 in Bladder Cancer. Cancers (Basel) 2024;16(6) doi 10.3390/cancers16061195.

11. Bharti R, Dey G, Lin F, Lathia J, Reizes O. CD55 in cancer: Complementing functions in a non-canonical manner. Cancer Lett 2022;551:215935 doi 10.1016/j.canlet.2022.215935.

12. Thongchot S, Jirapongwattana N, Luangwattananun P, Chiraphapphaiboon W, Chuangchot N, Sa-Nguanraksa D, et al. Adoptive Transfer of Anti-Nucleolin T Cells Combined with PD-L1 Inhibition against Triple-Negative Breast Cancer. Mol Cancer Ther 2022;21(5):727–39 doi 10.1158/1535-7163.MCT-21-0823.

13. Yu EY, Zhang H, Fu Y, Chen YT, Tang QY, Liu YX, et al. Integrative Multi-Omics Analysis for the Determination of Non-Muscle Invasive vs. Muscle Invasive Bladder Cancer: A Pilot Study. Curr Oncol 2022;29(8):5442–56 doi 10.3390/curroncol29080430.

14. Lodewijk I, Duenas M, Rubio C, Munera-Maravilla E, Segovia C, Bernardini A, et al. Liquid Biopsy Biomarkers in Bladder Cancer: A Current Need for Patient Diagnosis and Monitoring. Int J Mol Sci 2018;19(9) doi 10.3390/ijms19092514.

15. Nicolazzo C, de Berardinis E, Gazzaniga P. Liquid Biopsy for Predicting Bacillus Calmette-Guerin Unresponsiveness in Non-muscle-invasive Bladder Cancer. Eur Urol Oncol 2021;4(1):124–5 doi 10.1016/j.euo.2020.09.003.

16. Goncalves CM, Henriques SN, Santos RF, Carmo AM. CD6, a Rheostat-Type Signalosome That Tunes T Cell Activation. Front Immunol 2018;9:2994 doi 10.3389/fimmu.2018.02994.

17. Oliveira MI, Goncalves CM, Pinto M, Fabre S, Santos AM, Lee SF, et al. CD6 attenuates early and late signaling events, setting thresholds for T-cell activation. Eur J Immunol 2012;42(1):195–205 doi 10.1002/eji.201040528.

18. Orta-Mascaro M, Consuegra-Fernandez M, Carreras E, Roncagalli R, Carreras-Sureda A, Alvarez P, et al. CD6 modulates thymocyte selection and peripheral T cell homeostasis. J Exp Med 2016;213(8):1387–97 doi 10.1084/jem.20151785.

19. Henriques SN, Oliveira L, Santos RF, Carmo AM. CD6-mediated inhibition of T cell activation via modulation of Ras. Cell Commun Signal 2022;20(1):184 doi 10.1186/s12964-022-00998-x.

20. Gurrea-Rubio M, Fox DA. The dual role of CD6 as a therapeutic target in cancer and autoimmune disease. Front Med (Lausanne) 2022;9:1026521 doi 10.3389/fmed.2022.1026521.

21. Fox DA. The role of CD6 in autoimmune diseases. Cell Mol Immunol 2018;15(11):1001–2 doi 10.1038/s41423-018-0015-1.

22. Kenney HM, Rangel-Moreno J, Peng Y, Chen KL, Bruno J, Embong A, et al. Multi-omics analysis identifies IgG2b class-switching with ALCAM-CD6 co-stimulation in joint-draining lymph nodes during advanced inflammatory-erosive arthritis. Front Immunol 2023;14:1237498 doi 10.3389/fimmu.2023.1237498.

23. Chalmers SA, Ayilam Ramachandran R, Garcia SJ, Der E, Herlitz L, Ampudia J, et al. The CD6/ALCAM pathway promotes lupus nephritis via T cell-mediated responses. J Clin Invest 2022;132(1) doi 10.1172/JCI147334.

24. Ruth JH, Gurrea-Rubio M, Athukorala KS, Rasmussen SM, Weber DP, Randon PM, et al. CD6 is a target for cancer immunotherapy. JCI Insight 2021;6(5) doi 10.1172/jci.insight.145662.

25. Enyindah-Asonye G, Nwankwo A, Rahman MA, Hunegnaw R, Hogge C, Helmold Hait S, et al. Overexpression of CD6 and PD-1 Identifies Dysfunctional CD8(+) T-Cells During Chronic SIV Infection of Rhesus Macaques. Front Immunol 2019;10:3005 doi 10.3389/fimmu.2019.03005.

26. Pettenati C, Ingersoll MA. Mechanisms of BCG immunotherapy and its outlook for bladder cancer. Nat Rev Urol 2018;15(10):615–25 doi 10.1038/s41585-018-0055-4.

27. Zhang L, Luo L, Chen JY, Singh R, Baldwin WM, 3rd, Fox DA, et al. A CD6-targeted antibody-drug conjugate as a potential therapy for T cell-mediated disorders. JCI Insight 2023;8(23) doi 10.1172/jci.insight.172914.

28. Lin H, Zhou Q, Wu W, Ma Y. Midkine Is a Potential Urinary Biomarker for Non-Invasive Detection of Bladder Cancer with Microscopic Hematuria. Onco Targets Ther 2019;12:11765–75 doi 10.2147/OTT.S235134.

29. Vu Van D, Heberling U, Wirth MP, Fuessel S. Validation of the diagnostic utility of urinary midkine for the detection of bladder cancer. Oncol Lett 2016;12(5):3143–52 doi 10.3892/ol.2016.5040.

30. Wu K, Zeng J, Shi X, Xie J, Li Y, Zheng H, et al. Targeting TIGIT Inhibits Bladder Cancer Metastasis Through Suppressing IL-32. Front Pharmacol 2021;12:801493 doi 10.3389/fphar.2021.801493.

31. Hannouneh ZA, Hijazi A, Alsaleem AA, Hami S, Kheyrbek N, Tanous F, et al. Novel immunotherapeutic options for BCG-unresponsive high-risk non-muscle-invasive bladder cancer. Cancer Med 2023;12(24):21944–68 doi 10.1002/cam4.6768.

32. Fink EE, Sona S, Lee BH, Ting AH. Processing and cryopreservation of human ureter tissues for single-cell and spatial transcriptomics assays. STAR Protoc 2022;3(4):101854 doi 10.1016/j.xpro.2022.101854.

33. Hao Y, Hao S, Andersen-Nissen E, Mauck WM, 3rd, Zheng S, Butler A, et al. Integrated analysis of multimodal single-cell data. Cell 2021;184(13):3573–87 e29 doi 10.1016/j.cell.2021.04.048.

