## Supplementary figures and images for "Single Cell RNA-sequencing of BCG naïve and recurrent non-muscle invasive bladder cancer reveals a CD6/ALCAM-mediated immune-suppressive pathway"

### Supplemental Figures

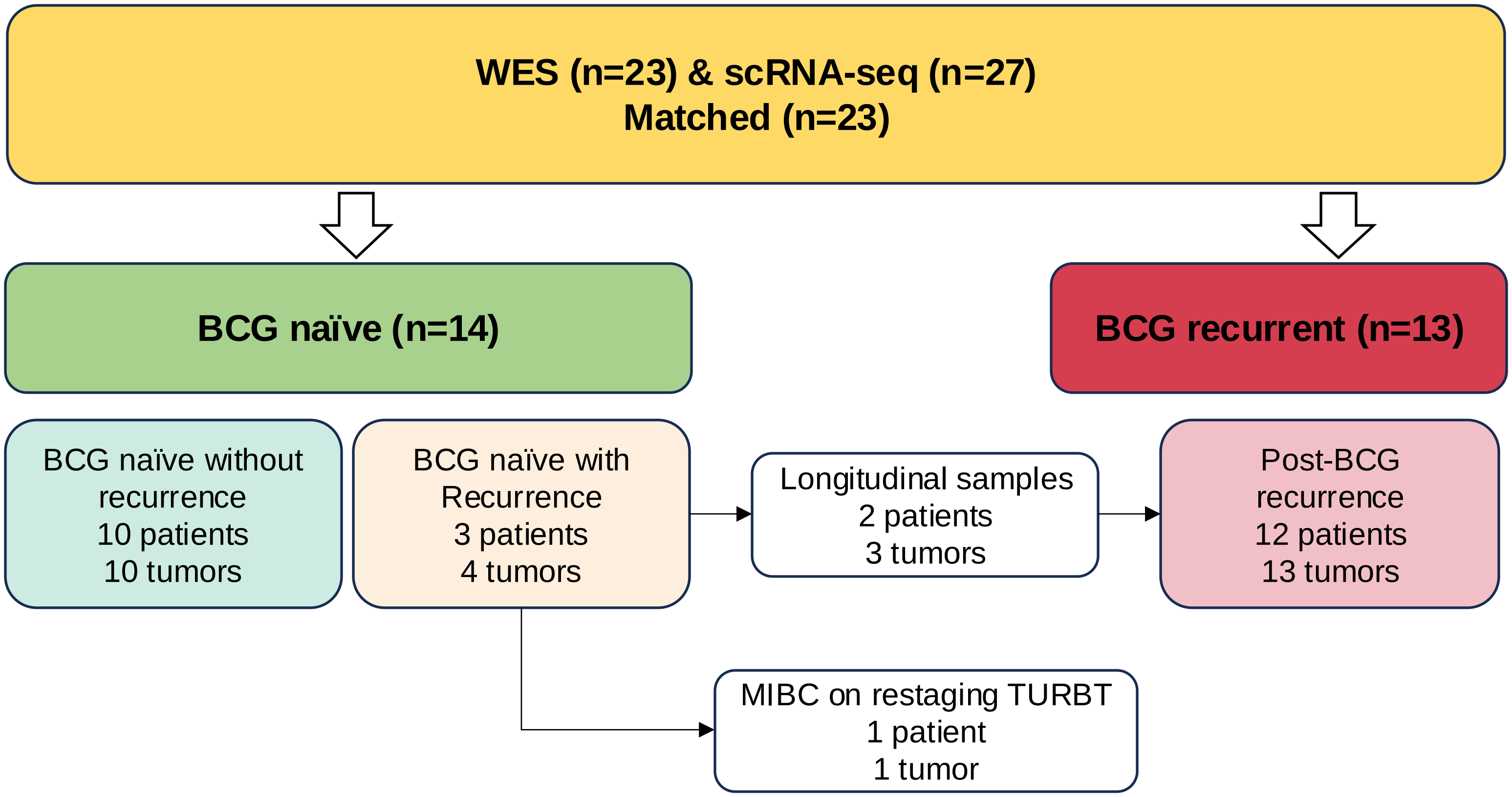

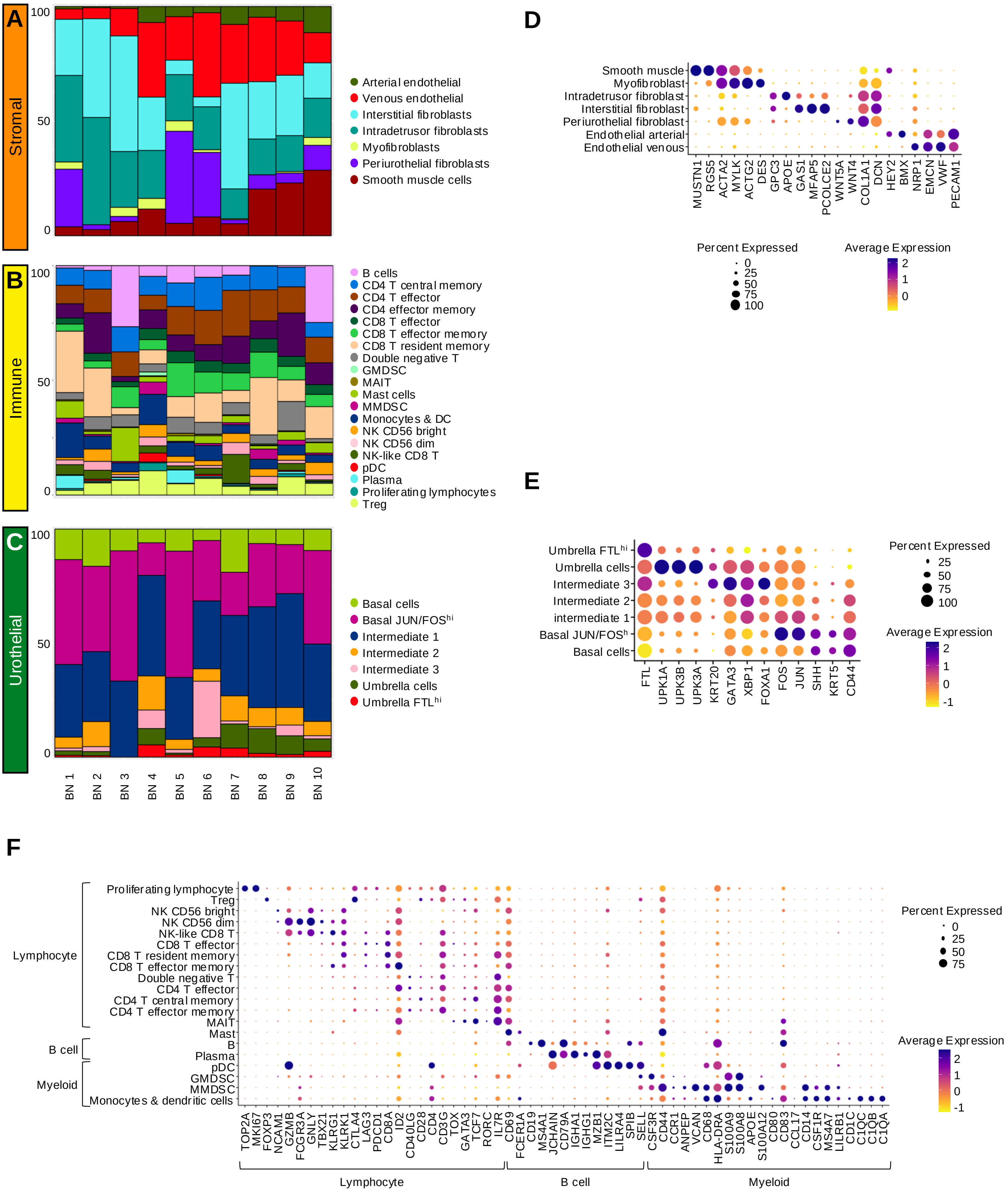

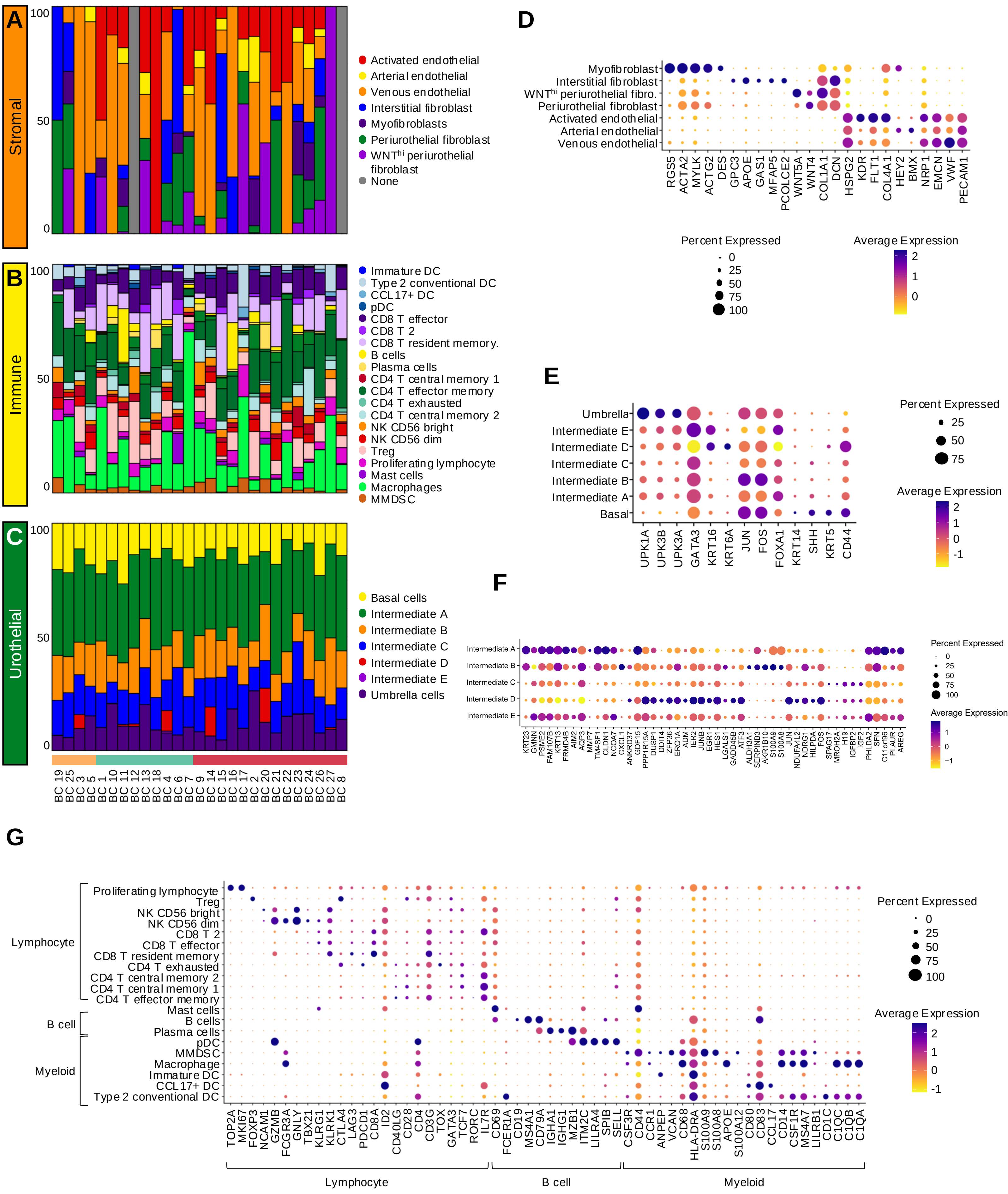

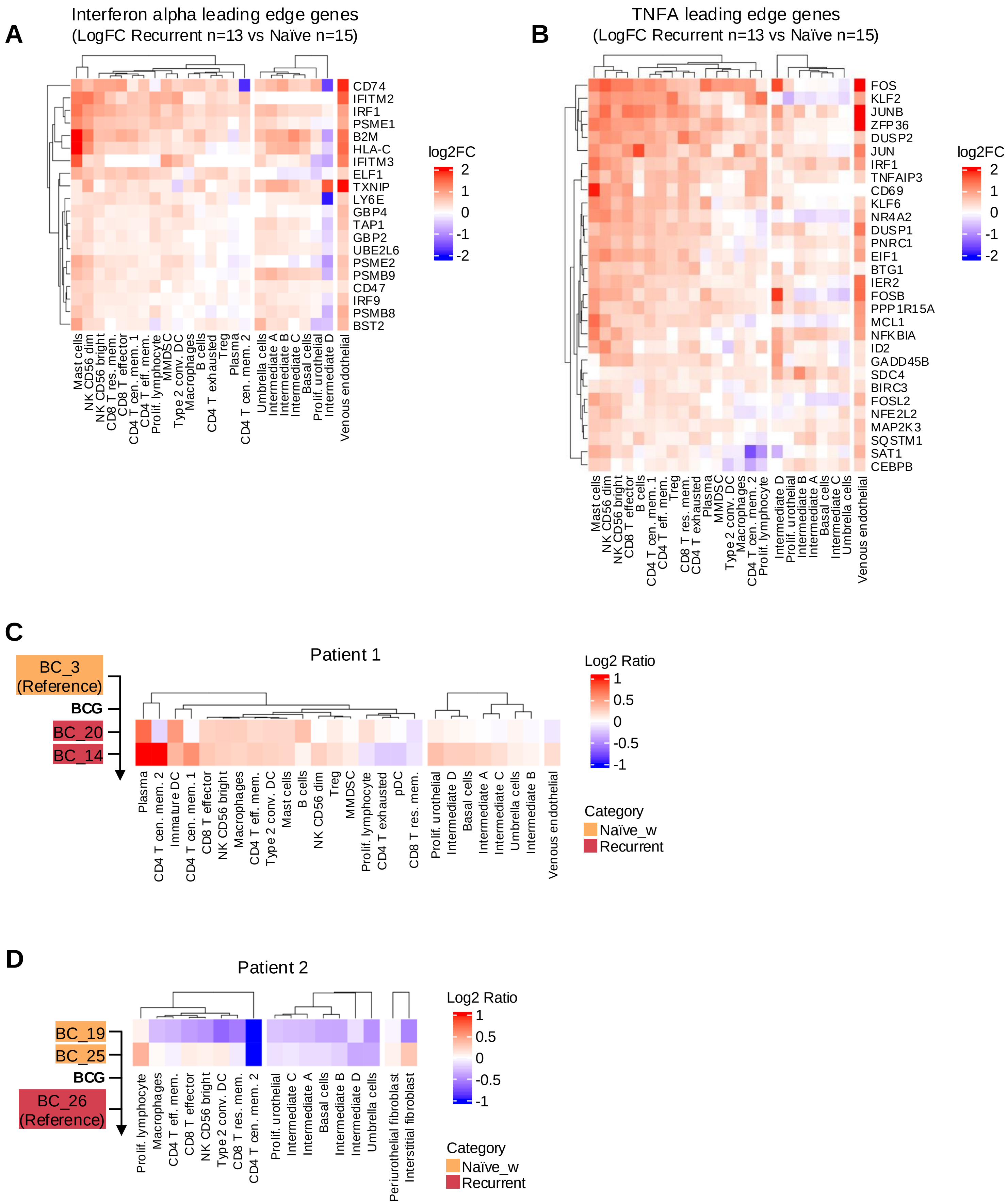

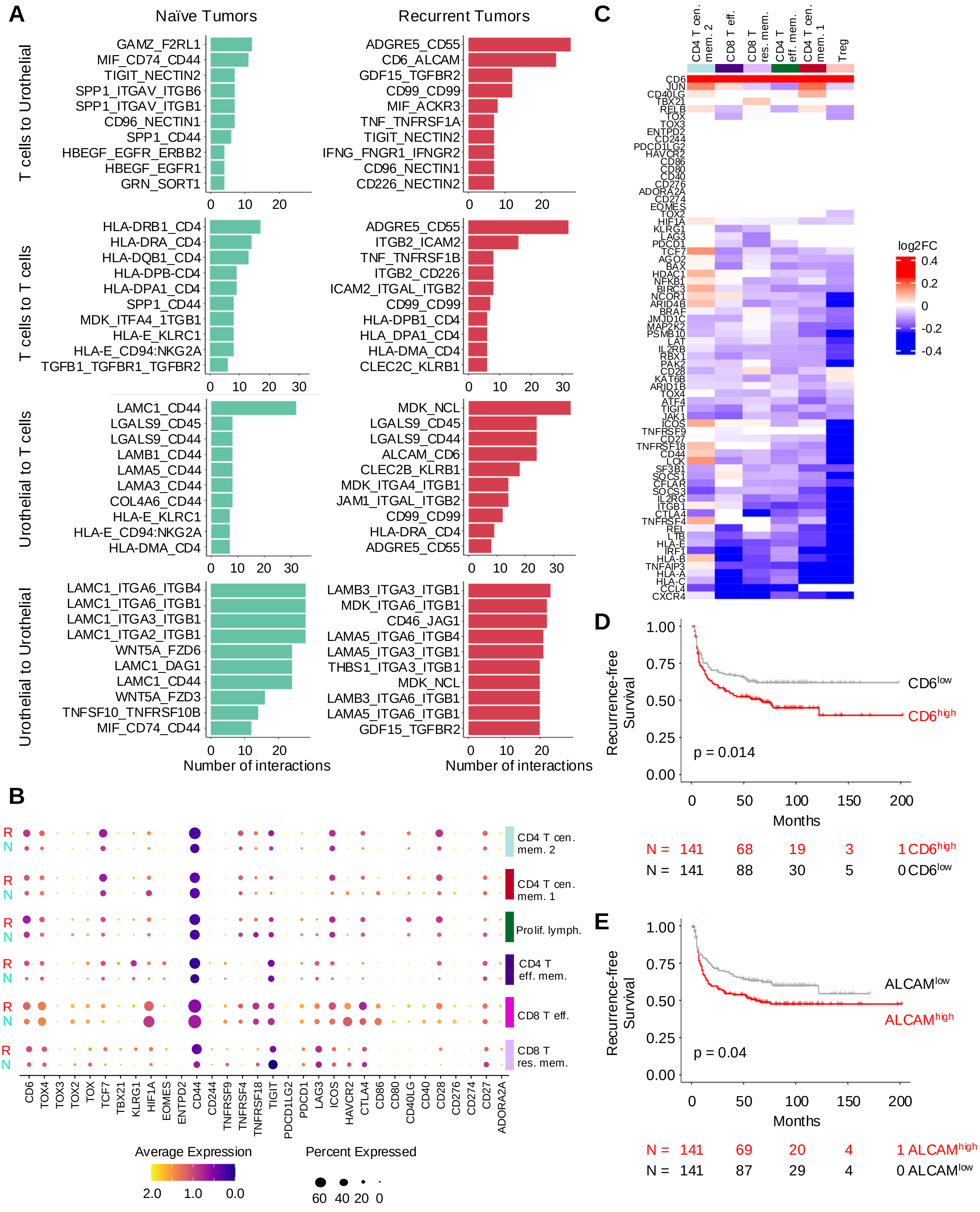
